# Detection of *Paracoccus yeei* in Spontaneous Bacterial Peritonitis using Rapid, Long Read Sequencing

**DOI:** 10.1101/2025.02.28.640566

**Authors:** Nasim Safaei, Zhi-Luo Deng, Valerie Ohlendorf, Beate Junk, Ludwig Sedlacek, Gemma L. Kay, Justin O’Grady, Jennifer Debarry, Etienne Ruppé, Markus Cornberg, Benjamin Maasoumy, Alice C. McHardy

## Abstract

Ascites is a common complication in patients with decompensated liver cirrhosis. Spontaneous bacterial peritonitis (SBP) is the most frequent infection, affecting up to 30% of hospitalized patients with ascites. Multidrug-resistant bacteria in patients with liver cirrhosis are becoming more common, particularly in nosocomial infections. With early detection and adequate antibiotic therapy, the mortality rate of SBP can be reduced from over 90% to 15-20%. However, the etiologic agent remains undetected by conventional pathogen diagnostics from ascites in more than half of the patients.

Here, we describe a robust workflow combining long read metagenome sequencing with best practices for low microbial biomass sample processing and bioinformatic analytics, to detect and characterize pathogens from the cirrhotic ascites of SBP patients, as a complement to routine microbiological diagnostics. This approach identified *Paracoccus yeei* as a likely etiologic agent for a patient who developed a fever after successful treatment of an *Escherichia coli* infection. Analysis of the genome of *P. yeei* recovered directly from ascites delineated pathogenicity-related features and an antimicrobial resistance profile consistent with the patient’s treatment. Pangenome phylogenetics placed the *P. yeei* strain closest to isolates associated with abdominal infections.

Our study underscores the utility of metagenomics for a rapid and comprehensive assessment of infectious agents, suggesting a novel and rarely reported pathogen contributing to SBP with potential implications for diagnostics and therapeutic strategies.

**Importance:** Our study demonstrates the clinical utility of real-time, long-read metagenomics in identifying elusive pathogens in Spontaneous Bacterial Peritonitis (SBP), a condition with high mortality and frequently inconclusive conventional diagnostics. By uncovering *Paracoccus yeei* as a likely secondary etiologic agent and characterizing its pathogenicity and AMR profile, our findings expand the known etiology of SBP and emphasize the potential of metagenomics to improve diagnostic accuracy, guide targeted treatment, and ultimately reduce mortality rates in infectious diseases. This approach could shape future clinical practice for managing complex infections.

## Introduction

Ascites is a common complication in patients with decompensated liver cirrhosis. Spontaneous bacterial peritonitis (SBP) is the most frequent infection in this group, affecting 10 to 30% of hospitalized patients with ascites(1). The prevalence of multidrug-resistant bacteria in patients with liver cirrhosis has been increasing over the last years and is particularly high in nosocomial infections. Without early detection and personalized adequate antibiotic therapy, the mortality rate of SBP is over 90%, though it can be substantially reduced to 15-20% by rapid diagnosis and targeted antibiotic treatment(2). However, the etiologic agent remains undetected by conventional cultivation-based pathogen diagnostics from ascitic fluid in more than 50% of patients(3). For SBP patients, the mortality rate is very high, at 32.5%(4) within a month of diagnosis, urgently requiring further improvements.

The emerging field of clinical metagenomics has the potential to substantially advance clinical infection research and diagnostics(5, 6). It enables rapid and comprehensive pathogen discovery directly from patient samples, the discovery of multiple pathogens(7), and substantially expands the range of discoverable taxa, to include those that are slow-growing, challenging to culture or not covered by targeted amplification assays(8). Moreover, the turnaround time of long-read sequencing of patient samples can be substantially reduced relative to cultivation-based diagnostics(6), which is crucial for the successful treatment of SBP. Furthermore, the approach is applicable to infectious disease diagnostics also in low and middle income countries(7). However, adapting clinical metagenomics to low microbial biomass clinical samples with suspected infections is challenging, requiring highly efficient removal of human host material and the ability to confidently distinguish between likely etiologic agents and low-abundant microbial contaminants from various sources(6, 9). Here, we demonstrate the value of an end-to-end analytic workflow using real-time, long read metagenome sequencing that we have established for detecting and characterizing the likely etiologic agents from the cirrhotic ascites of a patient diagnosed with spontaneous bacterial peritonitis.

## Materials and Methods

### Metagenomic sequencing

Metagenomic DNA extraction was performed utilizing the ZymoBIOMICS DNA/RNA Miniprep Kit from the patients’ ascites sample (case sample), from a frozen aliquot of the sample (case repeat), as a repetition, and Control_1 (water). For DNA extraction for Control_2 (water) the HostZERO™ Microbial DNA Kit was employed. To rule out batch-effects and potential reagent contamination from sequencing, case and no-template controls were sequenced in the same run and the case repeat was sequenced in another run without any further samples. Specifically, the case sample and no-template controls were sequenced together with two further samples using the Rapid PCR barcoding kit (SQK-RPB004) on the MinION sequencer for five hours, while the replicate case sample was sequenced without further samples using the Rapid PCR Barcoding Kit 24 V14 (SQK-RPB114.24) for 20 hours. Twenty-six ascites samples from individuals without SBP infection were also sequenced in different runs as negative biological cases, facilitating distinction between infection-related taxa and others introduced e.g. from other body sites by sampling. Host depletion, sample preparation and sequencing were performed as described in(6).

Due to shorter DNA fragment sizes and read lengths obtained in the repeated sequencing of frozen aliquot, additional false positive, low abundant taxa were identified in the case_repeat in comparison to the case with fresh sample material. Related assignments at genus level to *P. yeei* and *E. coli* likely also originate from these two taxa. *Moraxella osloensis* is a very common and abundant skin microbiome constituent(10) and contaminant in some samples.

### Taxonomic profiling

Minimap2 version 2.20(11) was used to remove human reads by mapping sequence data against the human reference genome version GRCh38. Subsequently, MetaX (version 0.8.0)(12) was used for taxonomic profiling, identifying taxa, genome coverage and their relative abundances in the patients’ sample. From the taxonomic profiles, phages and *Cutibacterium acnes* were excluded prior to genome coverage and relative abundance calculations. *Cutibacterium acnes* is the most abundant skin microbe, and present in all samples of the IHSMGC human skin microbiome dataset, with an average abundance of 28%(10), and was considered as a contaminant. From the obtained taxonomic profiles, we identified as likely etiologic agents those species present in both replicate metagenome sequence samples, but not in the controls (see Table S1 in the supplemental material). geNomad version 1.7.4(13) was used to identify mobile genetic elements including plasmids and viruses in the metagenomic data.

### Phylogenomic analyses

We pooled the two repeats of the metagenomic sequencing data from the patient sample then performed a metagenome assembly using metaFlye version 2.9.3(14). The contigs were taxonomically classified using Kraken2 version 2.1.3(15). For phylogenomic analyses of *Paracoccus* genomes, all *Paracoccus* genomes from NCBI RefSeq database with level “Complete” were downloaded. Prokka version 1.14.6(16) was used to annotate the genes in the genome assemblies. Panaroo version 1.5.0(17) was used to construct the pangenome for *Paracoccus* genomes. This analysis identified 28 conserved gene groups across the *Paracoccus* genomes. Then, a core gene alignment was built for these core gene groups. In the alignment, regions with gaps longer than 50 nucleotides were trimmed to prevent bias from low genome coverage of the clinical sample sequencing data and a phylogenetic tree was constructed using IQ-TREE version 1.6.12(18) with parameter setting “-m GTR+G -nt AUTO -bb 1000 -alrt 1000”. A second phylogenetic tree for *P. yeei*, was constructed by including in addition to *P. yeei* genome from the patients’ sample eleven *P. yeei* genomes available at RefSeq and the closely related *P. mutanolyticus,* using a core gene alignment of 174 gene groups. The *P. mutanolyticus* genome was used as an outgroup to determine the root of the tree. The alignment trimming and tree construction was done using the same workflow and parameter setting as for the *Paracoccus* genus tree above.

### Functional analyses

Taxonomic identifiers assigned to each metagenome sequence by MetaX were used for the genus-specific functional analysis. We conducted a functional annotation with pathogenicity categories, such as antibiotic resistance, induce inflammation, virulence regulator and secretion for the metagenomic sequences using SeqScreen version 4.3(19) with database SeqScreenDB_23.3. The Gene Ontology (GO) terms with assignment confidence score higher than 0.9 were used for the GO term enrichment analysis. We utilized ClusterProfiler version 4.12.0(20) to identify GO terms enriched in *P. yeei* compared to in other taxa. The “Benjamini-Hochberg” procedure was applied to adjust p-values to control for false discovery rate. GO terms with an adjusted p-value of less than 0.05 were considered as significantly enriched. The GO terms were visualized in two-dimensional semantic space by REVIGO(21) to reduce the redundancy introduced by the GO hierarchy, into three GO groups namely biological process (BP), molecular function (MF) and cellular component (CC). The uniprot hits with e-value below 1e-10 were kept to identify the sequences involved in microbial pathogenicity categories. The genus gene links were visualized with circlize(22) in the circular layout.

### Microbiological diagnostics

For microbiological diagnostics, ascitic fluid was either directly inoculated into blood culture bottles on the ward (BD Bactec), following the manufacturer’s instructions, and sent to the microbiology department, for an incubation period of seven days. When positive signals in blood culture bottles were detected, portions of the samples were placed on chocolate blood agar and columbia blood agar for the isolation of aerobic bacteria (48 hour incubation), and on Schaedler anaerobe agar (72h). Otherwise, if native ascitic fluid samples were sent to the microbiology department, incubation was performed on chocolate blood agar and Columbia blood agar (48 hour incubation), Schaedler anaerobe agar (96h) and anaerobic Sodium thioglycolate broth (7 days) for anaerobic microorganisms. If growth was detected, pathogens were identified by standard microbiology methods, e.g. utilization of Matrix-Assisted Laser Desorption/Ionization Time-of-Flight (MALDI-TOF) mass spectrometry.

### Ethics approval

The prospective registry (INFEKTA) for infectious complications in liver cirrhosis and ascites is registered as DRKS00010664 (German Register for Clinical Trials). All participants provided written informed consent before participation in the study, as required by the Hannover Medical School Ethics Committee (Approval No. 3188-2016).

## Results

### Sample and patient description

A 38-year-old patient with decompensated liver cirrhosis, classified as Child-Pugh stage B, with Primary Sclerosing Cholangitis as the underlying liver disease, was urgently admitted to the hospital in June 2023. The patient was in a deteriorating general condition and reported increasing amounts of ascites. Recently, ascites punctures had been required at least once a month. In April 2023 a prior episode of SBP had been treated according to an established antibiogram for *Escherichia coli* with piperacillin/tazobactam and he had since been placed on norfloxacin as prophylaxis. The patient also had dialysis-dependent kidney insufficiency related to type 3 diabetes mellitus and received immunosuppressive medication because of a prior lung transplantation due to cystic fibrosis. Upon admission, laboratory results revealed substantially elevated blood levels of C-reactive protein (CRP) (229.6 mg/l), indicative of severe inflammation, Procalcitonin (PCT, 1.3 µg/l) and elevated ascites levels of polymorphonuclear leukocytes (2437 PMN/µl)(23), suggesting a recurrent episode of SBP (see Table S2 in the supplemental material). Consequently, treatment with piperacillin/tazobactam was initiated. On day 2, blood and ascites samples were sent for microbiological diagnostics. Ascites samples from the following day were subjected to metagenomic shotgun sequencing (Fig. 1a). To ensure reproducibility, a repeat case sample was sequenced in a separate run from the original case sample. Additionally, 26 samples from individuals without SBP infection were sequenced as negative cases, to distinguish between infection related taxa and other taxa, e.g. taxa potentially introduced by sampling from other body sites. Two no-template controls were included to monitor for potential contamination from the reagents.

**Fig. 1.**
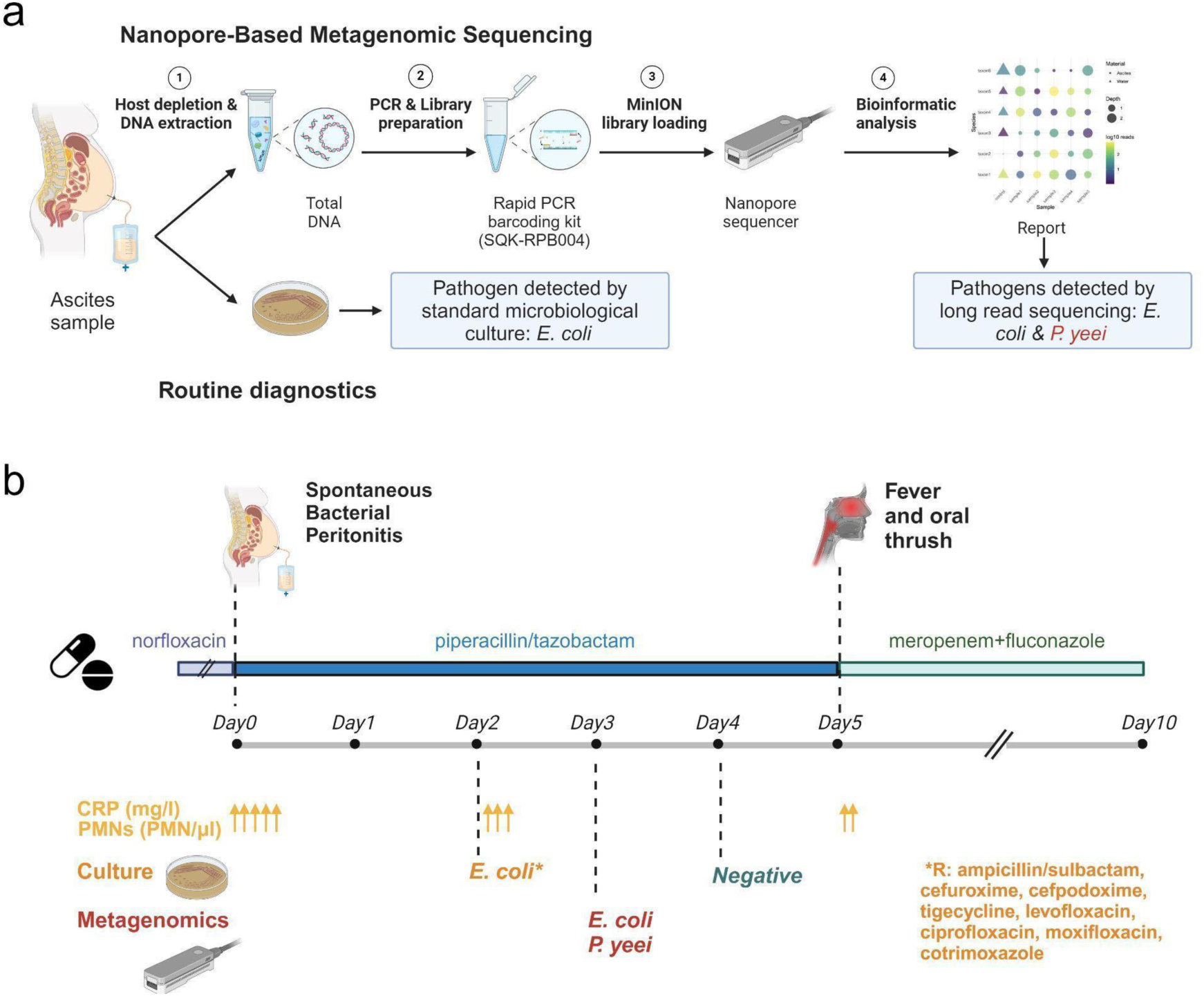
Microbial pathogen diagnostics using clinical metagenomics. (a) Steps in nanopore-based metagenomic sequencing vs. cultivation-based approach in routine diagnostics to identify pathogens from a patient sample. (b) Clinical timeline of the patient diagnosed with spontaneous bacterial peritonitis, detailing the antibiotic treatments administered and the diagnostic outcomes from both culture and metagenomics. Monitored CRP and PMN levels as inflammation markers (see Table S2 in the supplemental material) are indicated by arrows. Days after admission indicate the days of sample collection. Cultivation of the patients’ ascites samples detected *E. coli*, while metagenomic analysis identified both *E. coli* and *P. yeei*. The asterisk indicates the antibiotic resistance profile of *E. coli* determined in routine diagnostics. This figure was created with BioRender.com.

### Long read metagenomics identifies a second pathogen

Both microbiological diagnostics and metagenomic analysis revealed the presence of *E. coli* in the patients’ ascites: Microbiological diagnostics of a sample from two days after admission identified *E. coli* as a likely pathogen. The antibiogram indicated susceptibility of the pathogen to meropenem and piperacillin/tazobactam, together with a resistance to the fluoroquinolone moxifloxacin (see Fig. 1b and Table S3 in the supplemental material), which the patient had been given as a prophylaxis.

Three days after the admission, *P. yeei* was identified as a likely second pathogen by metagenomic analysis, with more than 10-fold higher relative abundance than *E. coli* in ascites sample, with 524 reads obtained after five hours of sequencing (see Fig. 2a and Table S1 in the supplemental material). To comply with suggested best practices for metagenome sequencing of samples with low microbial biomass(9) for investigating biological or technical contamination as an alternative source of the organism, we included multiple no-template controls in the sequencing experiments, performed a replicate metagenome sequencing of the patients’ sample and 26 ascites samples from SBP-negative patients, which uniquely identified *P. yeei* in the replicate patient samples of the patient with SBP. Additionally, several fragments of *Rhizobium pusense*, which is also an opportunistic, human pathogen(24), *Pseudomonas* and *Corynebacterium* were found exclusively in the patients’ ascites metagenome replicates. However, the substantially lower read numbers for these taxa provide less support for their clinical relevance than for *P. yeei.* Furthermore, we studied 26 additional ascites samples from patients without SBP using long-read metagenomics. Except for one, none of these included reads of *P. yeei.* The exception was a sample from another patient with abdominal paracentesis, where very few (200-fold less) *P. yeei* reads were found, likely resulting from contamination via the sampling site.

**Fig. 2.**
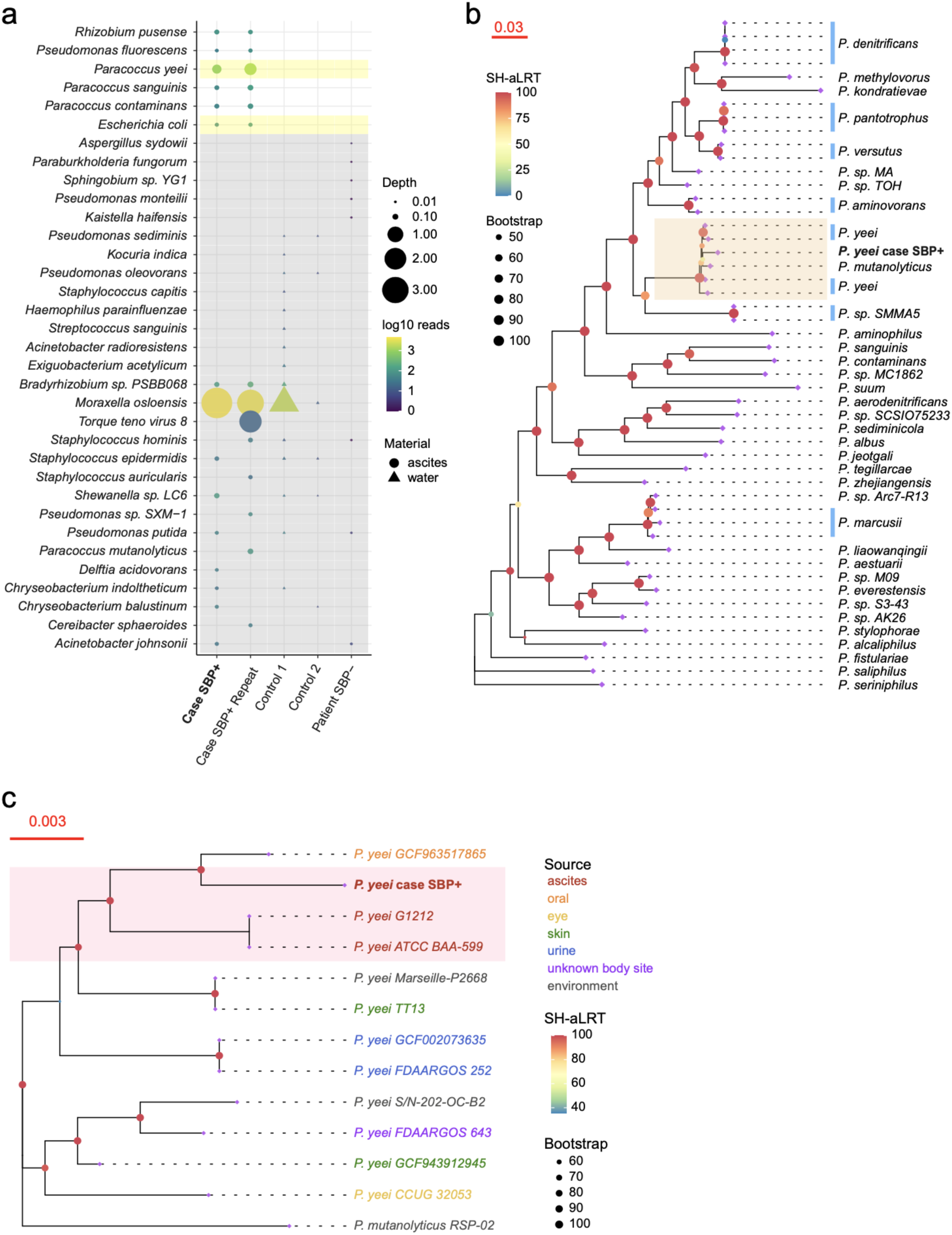
Taxonomic composition of patient and control samples and pangenome phylogeny of *Paracoccus* genomes. (a) Microbial species identified from metagenome sequencing data of the patients’ ascites sample (case, repeat_case) and no-template controls and the latest sample from an SBP-negative patient are shown (Materials and Methods). Point sizes are proportional to the depth of genome coverage for individual taxa estimated by MetaX, and the color indicates the log10 transformed read counts.To facilitate the overview, only up to 15 species with more than five reads are shown for each sample, except for the ascites sample of the patient without SBP, which had very few reads and thus all taxa are shown. Yellow bars indicate the likely etiological agents as those taxa that are present in the case samples, but not in the controls, with either the highest abundance (*P. yeei*) or confirmed by culture in routine diagnostics (*E. coli*). The gray shades depict all the taxa not present in both case samples and thus likely false positives. (b) Phylogenetic pangenome analysis of *Paracoccus* genomes from RefSeq and *P. yeei* genome from the patient’s metagenomic sample. The adjacent genomes on the tree from the same species are marked in light blue vertical bars. The branches shaded in yellow are the *P. yeei* and *P. mutanolyticus* genomes. (c) A phylogenetic tree comprising all *P. yeei* and *P. mutanolyticus* genomes from RefSeq, alongside the *P. yeei* genome assembled from the patient’s sample. The tip lab text color indicates the source of the genome being sampled or isolated. The pink-shaded branches represent genomes derived from human ascites samples. In (b) and (c) the color of the internal nodes indicates the SH-aLRT support value from IQ-TREE, and their size reflects the Bootstrap support value.

While infection parameters initially improved, and new blood and ascitic fluid cultures on day four post-admission yielded negative results (Fig. 1b), indicating successful treatment of the identified *E. coli*, the patient developed a fever of 38.3°C on the fifth-day post-admission. Furthermore, he presented symptoms of a sore throat (*Candida albicans* cultured from a throat swab) and oral thrush, prompting the addition of the antimycotic fluconazole to the treatment. As the patient also encountered frequent episodes of loose bowel movements, comprehensive stool analyses with multiplex PCR were performed, but revealed negative results for Rotavirus antigen, Norovirus, Adenovirus, and Cytomegalovirus (CMV) DNA. Also, no pathogenic intestinal bacteria, including *Clostridioides difficile* were detected. Because of the fever and continuously elevated CRP, Meropenem was prescribed for the next five days, after which infection markers improved.

### Phylogenomic analysis of *Paracoccus yeei*

To characterize the evolutionary provenance of the *P. yeei* genome that was recovered from the patients’ ascites metagenome, we performed a pangenome phylogenomic analysis. To this end, conserved marker genes were identified for all available genomes of the *Paracoccus* clade and used to reconstruct evolutionary relationships via maximum likelihood tree inference. The *P. yeei* genome from the patients’ ascites forms a stable sublineage with the other strains of *P. yeei* (Fig. 2b). Given *P. yeei’s* presence in diverse environmental settings such as soil and water(25, 26), we sought to determine if the strain from our sample is clinically relevant rather than merely an environmental contaminant. Next, we generated another phylogenetic tree including all *P. yeei* genomes from the RefSeq database of both clinical and environmental samples and *P. mutanolyticus*. In this analysis, the *P. yeei* from the patients’ ascites was most closely related to a *P. yeei* metagenome-assembled genome (MAG, GCF_963517865) from the human oral microbiota(27) and as well as a *P. yeei* isolate genome (G1212) from abdominal dialysate(28) (Fig. 2c), with more divergence relative to the ten available other strains.

### Functional analysis of the *P. yeei* genome sequences

To elucidate the functional potential of the *P. yeei* genome in the patients’ ascites, we performed a Gene Ontology (GO) term enrichment analysis (Fig. 3a-c, Materials and Methods), characterizing the functional groups substantially enriched in *P. yeei* sequences relative to other sequences in the metagenome sample. This indicated a pronounced overrepresentation of various functional groups associated with pathogenicity. These include response to antibiotics, SOS response, protein repair, bacterial-type flagellum, toxin activity, and antioxidant activity. Additionally, categories related to DNA transposition, DNA integration, outer membrane, chemotaxis, extracellular polysaccharide biosynthetic processes, type I secretion system, proteinase/peptidase activities, hydrolase activity, and amino acid transport are also notably enriched. Further enriched functions encompass ABC transporter systems, xenobiotic transport, actin binding, and iron binding.

**Fig. 3.**
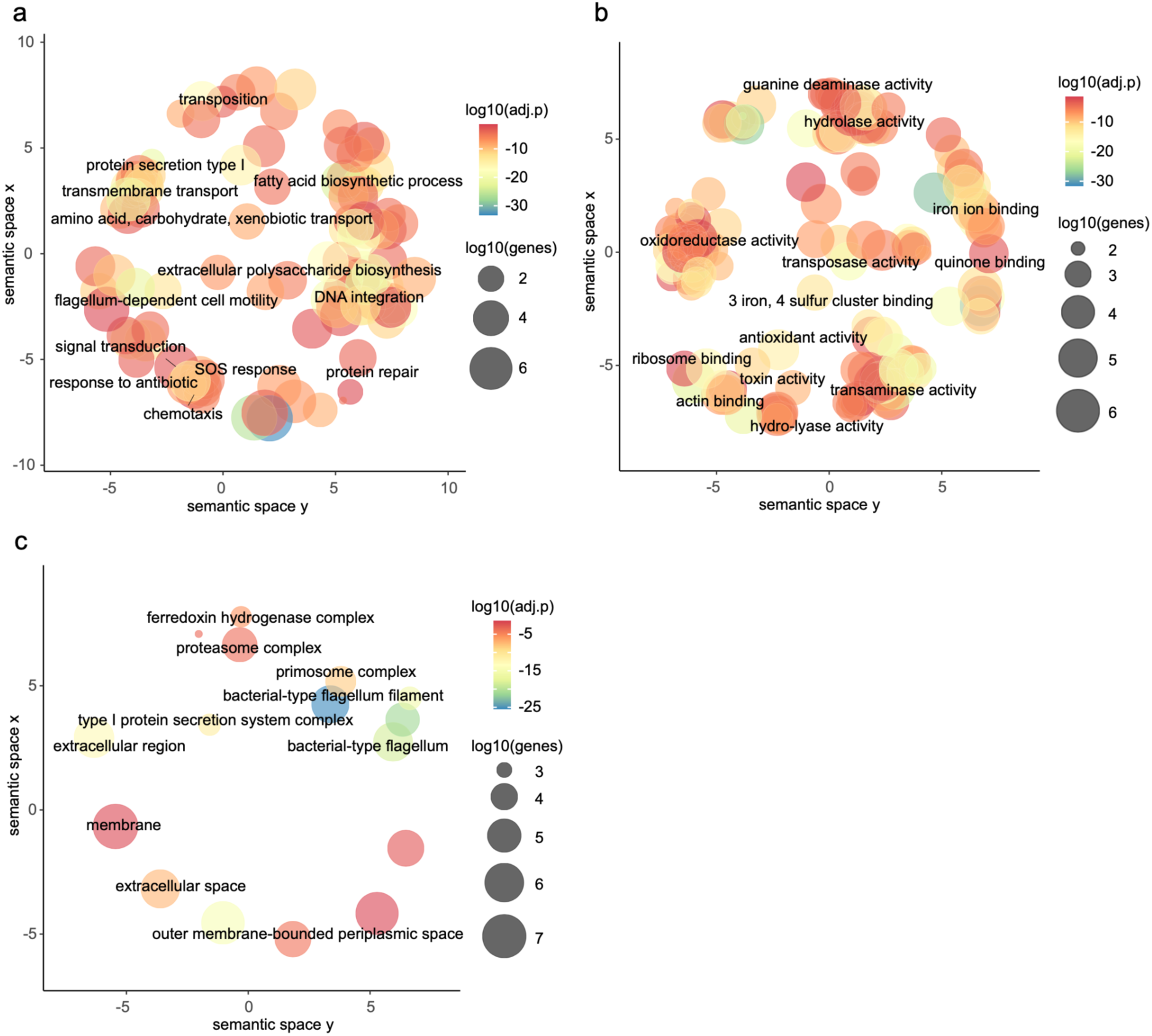
Semantic similarity of GO terms enriched for the metagenomic *P. yeei* sequences relative to other metagenomic data, categorized into (a) biological process (BP), (b) molecular function (MF), and (c) cellular component (CC) The color scale from red to blue indicates the significance of enrichment, with blue being the most strongly enriched. Circle size represents the number of sequences associated with that GO term in the metagenomic data. A larger circle indicates a more general term.

### Antimicrobial resistance and other pathogenicity genes

We further characterized genes implied in pathogenicity and related processes from *Paracoccus* and other genera from the patients’ metagenomes (Fig. 4, Materials and Methods). Of the 21 gene sequences found for *Paracoccus*, 15 are implicated in processes relating to antimicrobial resistance (AMR), four encode *RbsA* involved in transmembrane transport and secretion, and two are associated with inflammation (see Fig. 4a and Table S4 in the supplemental material). Eight secretion-related gene sequences and two AMR gene sequences were identified for *Rhizobium*, for which genomic data were also found exclusively in the patients’ sample, but not in controls (Fig. 2a). Among the 15 AMR-associated sequences identified in *Paracoccus*, two sequences encode genes annotated with the multidrug efflux pump gene *emrB*. One sequence encodes a gene conferring resistance to beta-lactams, ten sequences are associated with resistance to Bacitracin, and two sequences are related to Fosfomycin resistance (Fig. 4b). The multidrug efflux pump protein EmrB, a member of the MFS efflux pump family, has been demonstrated to confer resistance against fluoroquinolones(29–33), beta-lactams, and tazobactam in various clinically relevant bacterial species. The identified beta-lactam resistance gene is homologous to an HcpB-like beta-lactamase, exhibiting moderate hydrolytic activity(34). A closer examination of the taxonomy placed these AMR associated sequences with *P. yeei*. As mobile genetic elements oftentimes confer resistance to antibiotics, but cannot be easily assigned taxonomically in case of broader host ranges, we analyzed the AMR genes in the taxonomically unclassified sequences. This identified another five sequences encoding beta-lactamase genes and another gene encoding EmrB (Fig. 4c), aligning in general with the patients’ treatments with beta-lactam antibiotics (Fig. 1). One of these which has length of ∼3 kb was predicted to be plasmid-derived. The first ⅔ is homologous to a wide range of taxa including Paracoccaceae (with 92.11% identity), whereas the remainder matches a type A beta-lactamase. As *P. yeei* strains typically harbor multiple plasmids and/or extrachromosomal replicons(35), it is plausible that this putative beta-lactamase originates from a *P. yeei* plasmid. Taken together, the metagenome analysis suggests that resistance-conferring genes against the beta-lactam antibiotics (piperacillin/tazobactam) that were administered over the first five days of the patients’ hospital stay accumulated in the metagenome, including the reconstructed *P. yeei* genome.

**Fig. 4.**
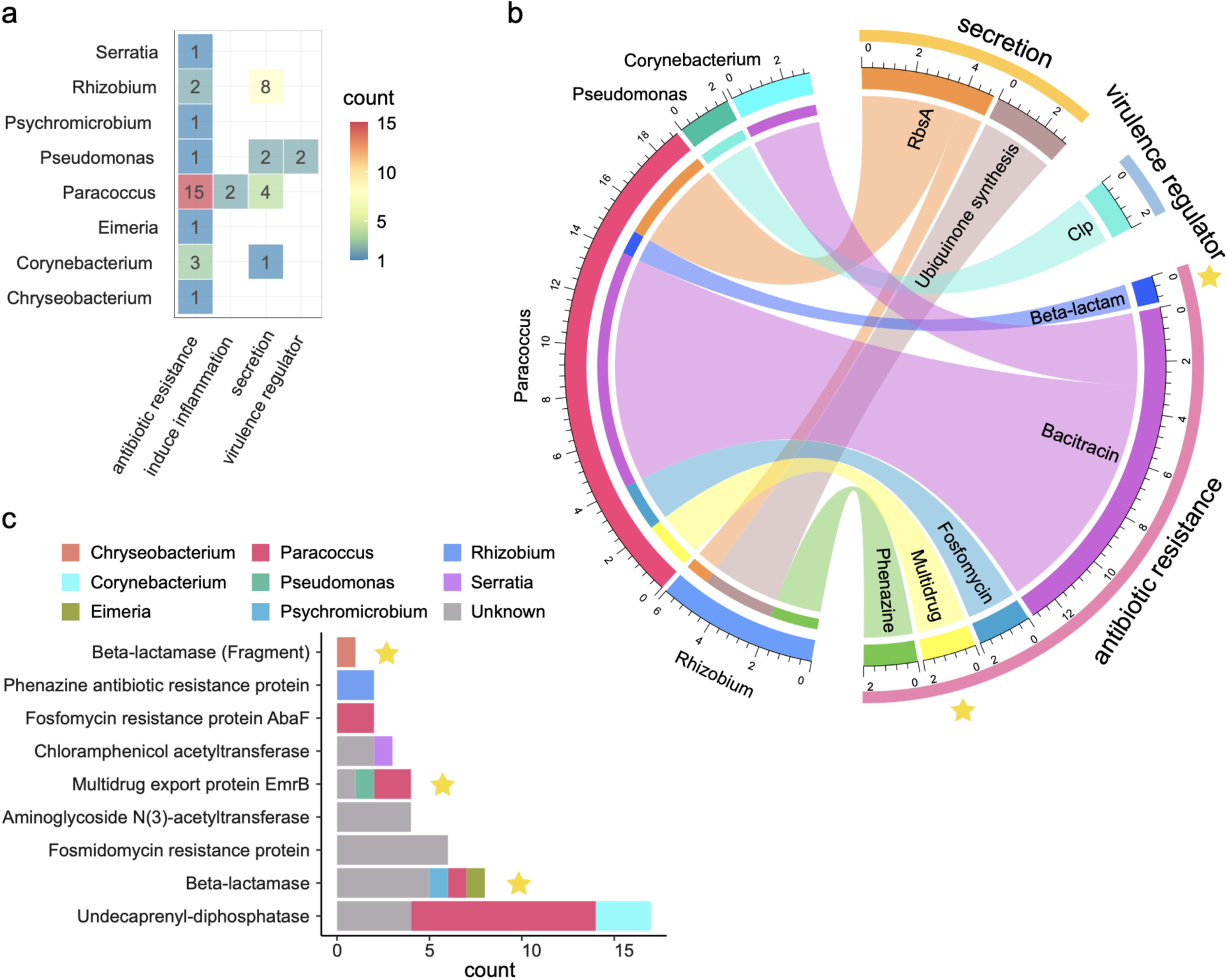
Microbial pathogenicity genes in the patients’ ascites metagenome. The number of sequences from different genera involved in pathogenicity categories (a). The sequences encoding specific pathogenicity genes in different genera (b). The number of sequences from different genera and unclassified sequences that encode different AMR genes. Genes highlighted by a yellow star represent AMR genes, which contribute to resistance for the fluoroquinolone and beta-lactam antibiotics administered to the patient.

## Discussion

Here, we describe how rapid metagenomic sequencing and analysis of an ascites sample from a patient with SBP led to the identification of two pathogens. One of these, *E. coli*, was confirmed by routine diagnostics, while the second one, *P. yeei*, was not. *P. yeei* is a Gram-negative bacillus from the Pseudomonadota (formerly Proteobacteria) phylum and is recognized as an opportunistic pathogen in humans. It is associated with various clinical conditions, including peritonitis. Diagnosing *P. yeei* in microbiological cultures poses challenges, however, as an initial Gram stain may not reveal the presence of the organism(36, 37), and it requires specialized media and extended incubation for growth(38, 39) not routinely employed in microbiological diagnostics or may be overgrown by other microbes.

While successful treatment of the *E. coli* with the initially administered beta-lactam antibiotics was confirmed by a negative result from cultivation-based diagnostics two days later, the patients’ continuously elevated infection markers suggested the presence of another, undetected pathogen, and prompted the prescription of meropenem as an alternative antibiotic. The metagenomic analysis of the patients’ ascites sample under the treatment with piperacillin/tazobactam identified *P. yeei* present at 10-fold higher abundance than remnants of *E. coli* and several other low-abundant taxa. As ascites samples from patients with peritonitis have low (or no) microbial biomass, distinguishing between truly present taxa representing likely etiologic agents and biological and technical contaminants is required. In line with suggested best practices(9), we sequenced multiple types of negative controls and technical sample replicates, which allowed us to identify multiple taxa as likely technical contaminants. To detect typical biological contaminants, e.g. from the sampling site, we screened 26 additional ascites samples from patients without SBP using metagenomic sequencing. However, *P. yeei* was not detected at notable abundances in any other samples.

*P. yeei* has been previously identified as a pathogen in peritonitis, particularly in patients with diabetes (see Table S5 in the supplemental material)(38–41), as the patient studied here. Consistent with it being a second relevant pathogen, in pangenomic analysis, the *P. yeei* strain from this study aligned phylogenetically closer to strains found in abdominal dialysate of an infected patient than to strains from the environment and skin. Diabetes mellitus often leads to immune system dysfunction(42) and frequently progresses to renal failure, necessitating peritoneal dialysis. Both peritoneal dialysis, as well as peritoneal paracentesis significantly increase the risk of the entry of environmental pathogens such as *P. yeei*(37). The combination of diabetes-induced immunosuppression and the procedural risks associated with peritoneal dialysis enhance the susceptibility to peritonitis with *P. yeei*(*25, 39*). Our findings suggest that similarly patients with peritoneal paracentesis and underlying conditions such as diabetes, might have to be carefully managed to prevent such infections.

*P. yeei* oftentimes is susceptible to beta-lactams, fluoroquinolones, aminoglycosides, and carbapenems(25), though resistances to some beta-lactams and fluoroquinolones are described(35, 38, 41). This is consistent with the metagenomic analysis of AMR genes, which identified a gene in *P. yeei* encoding an HcpB-like beta-lactamase. Furthermore, an *emrB*-like gene was identified in the plasmid sequences of *P. yeei* as a major component of a multidrug efflux pump system conferring AMR against multiple antibiotics including fluoroquinolones, and multiple taxonomically unclassified sequences were found to encode beta-lactamase genes. These sequences may belong to mobile genetic elements, thereby contributing to the AMR. These findings indicate resistance patterns to the administered piperacillin/tazobactam therapy, and aligns with the observed patient recovery after switching to the carbapenem antibiotic meropenem.

Remarkably, functional analysis of the metagenomic data revealed various pathogenicity-related functional groups overrepresented in the *P. yeei* genome that was recovered from the ascites metagenome. These enriched categories suggest that it employs a suite of strategies to interact with and adapt to the host environment for establishing persistent infections. Key mechanisms include host cell invasion, nutrient sequestration, toxin secretion, eliciting inflammatory responses, immune evasion, antibiotic resistance, and biofilm formation. The enrichment of functional groups associated with DNA transposition and integration indicates *P. yeei* adapts to stressful conditions, such as those induced by antimicrobial treatment, by acquiring mobile genetic elements from its environment and surrounding cells(43, 44). Additionally, compared to other taxa in the sample, the *P. yeei* strain shows an overrepresentation of the SOS response and protein repair mechanisms. These processes are crucial for mitigating DNA damage and protein misfolding often induced by antibiotics such as fluoroquinolones, beta-lactams, and aminoglycosides(45, 46). Notably, the patient did not respond well to the administered antibiotics from the first two classes, fluoroquinolones and beta-lactams, during the treatment.

These findings highlight the potential of *Paracoccus* to contribute to pathogenic processes within clinical settings, warranting further investigation into its role in SBP etiology. Several secretion and AMR gene-related sequences were also characterized in *R. pusense*, despite its relatively lower abundance compared to *P. yeei*, supporting the hypothesis of a potential clinical relevance in infectious diseases(24).

## Data Availability

The sequencing data generated in this study can be accessed via accession PRJEB68145 from the European Nucleotide Archive. The MetaX taxonomy profiles for negative controls are available at https://doi.org/10.5281/zenodo.13829418.

## Acknowledgements

This work was supported by the German Center for Infection Research [TI 12.002]. Additionally, this work was partially funded by the German Research Foundation (DFG), through the PRACTIS–Clinician Scientist Program of Hannover Medical School [ME 3696/3-1] and the Cluster of Excellence RESIST [EXC 2155] of the German Research Foundation [Project number 390874280].

We thank Jörn Kalinowski and Tobias Busche of the Department of Genetics, Faculty of Biology, University of Bielefeld, Bielefeld and the Technology Platform Genomics, Center for Biotechnology, Bielefeld University for providing training and advise on Minion sequencing, Cordula Hege and Carmen Paulmann for editing the manuscript and the Center for Individualized Infection Medicine for providing lab space and equipment for the project.

