## Supplementary tables for "Detection of *Paracoccus yeei* in Spontaneous Bacterial Peritonitis using Rapid, Long Read Sequencing"

**S1 Table. Taxonomic profile at the species level identified with long lead metagenome sequencing.** The patients' ascites sample, two non-template controls, one SBP- case (Methods). Depth specifies the genome coverage for individual taxa estimated by MetaX, and NumReads the number of sequence fragments assigned to them. See Fig. 2 for data visualization.

| Case SBP+ | | | Case SBP+ Repeat | | | Control 1 | | | Control 2 | | | Patient SPB- | | |
|---|---|---|---|---|---|---|---|---|---|---|---|---|---|---|
| **Species** | **Depth** | **Num Reads** | **Species** | **Depth** | **Num Reads** | **Species** | **Depth** | **Num Reads** | **Species** | **Depth** | **Num Reads** | **Species** | **Depth** | **Num Reads** |
| *Moraxella osloensis** | 3.92 | 4,754 | *Moraxella osloensis** | 2.90 | 4,200 | *Moraxella osloensis** | 2.17 | 2,179 | *Moraxella osloensis** | 0.01 | 13 | *Pseudomonas putida* | 0.001 | 4 |
| *Paracoccus yeei* | 0.25 | 524 | *Torque teno virus 8* | 1.90 | 17 | *Bradyrhizobium sp. PSBB068* | 0.04 | 192 | *Staphylococcus epidermidis* | 0.01 | 8 | *Staphylococcus hominis* | 0.0007 | 1 |
| *Shewanella sp. LC6* | 0.05 | 129 | *Paracoccus yeei* | 0.48 | 1,162 | *Staphylococcus epidermidis* | 0.02 | 22 | *Pseudomonas oleovorans* | 0.004 | 6 | *Kaistella haifensis* | 0.0005 | 1 |
| *Bradyrhizobium sp. PSBB068* | 0.04 | 142 | *Bradyrhizobium sp. PSBB068* | 0.07 | 257 | *Exiguobacterium acetylicum* | 0.02 | 31 | *Pseudomonas sediminis* | 0.003 | 7 | *Pseudomonas monteilii* | 0.0003 | 1 |
| *Rhizobium pusense* | 0.04 | 86 | *Paracoccus sanguinis* | 0.06 | 126 | *Staphylococcus hominis* | 0.01 | 12 | *Chryseobacterium balustinum* | 0.002 | 6 | *Sphingobium sp. YG1* | 0.0002 | 1 |
| *Paracoccus contaminans* | 0.04 | 49 | *Paracoccus mutanolyticus* | 0.06 | 115 | *Acinetobacter radioresistens* | 0.01 | 22 | *Shewanella sp. LC6* | 0.002 | 6 | *Paraburkholderia fungorum* | 0.0001 | 1 |
| *Paracoccus sanguinis* | 0.04 | 57 | *Paracoccus contaminans* | 0.06 | 85 | *Streptococcus sanguinis* | 0.01 | 13 | | | | *Aspergillus sydowii* | 0.0001 | 2 |
| *Staphylococcus epidermidis* | 0.03 | 30 | *Staphylococcus hominis* | 0.03 | 40 | *Pseudomonas putida* | 0.01 | 46 | | | | | | |
| *Acinetobacter johnsonii* | 0.02 | 29 | *Rhizobium pusense* | 0.03 | 94 | *Haemophilus parainfluenzae* | 0.01 | 11 | | | | | | |
| *Escherichia coli* | 0.02 | 54 | *Staphylococcus auricularis* | 0.02 | 22 | *Chryseobacterium indoltheticum* | 0.01 | 21 | | | | | | |
| *Chryseobacterium balustinum* | 0.02 | 42 | *Pseudomonas fluorescens* | 0.02 | 71 | *Shewanella sp. LC6* | 0.01 | 27 | | | | | | |
| *Pseudomonas putida* | 0.02 | 49 | *Pseudomonas sp. SXM-1* | 0.02 | 70 | *Staphylococcus capitis* | 0.01 | 8 | | | | | | |
| *Pseudomonas fluorescens* | 0.01 | 37 | *Escherichia coli* | 0.02 | 83 | *Pseudomonas oleovorans* | 0.01 | 19 | | | | | | |
| *Chryseobacterium indoltheticum* | 0.01 | 24 | *Cereibacter sphaeroides* | 0.02 | 39 | *Kocuria indica* | 0.01 | 11 | | | | | | |
| *Delftia acidovorans* | 0.01 | 39 | | | | *Pseudomonas sediminis* | 0.01 | 12 | | | | | | |

**Notes:**

SBP: Spontaneous bacterial peritonitis.

*The bacterium *Moraxella osloensis* was identified as the most abundant species in the sample. Given its ubiquity and abundance in the skin microbiota (present in 99.9% samples of the IHSMGC dataset with an average abundance of 5.4%), it was considered as a contamination and disregarded.

**S2 Table. Dynamic clinical parameters for the SBP patient within the first five days after admission**

| Indicators | On admission day | Two days after | Five days after |
|---|---|---|---|
| CRP(mg/l) | 229.6 | 152.0 | 104.6 |
| PCT (µg/l) | 1.3 | .. | .. |
| AST(U/l) | 38 | 23 | 37 |
| ALT(U/l) | 24 | 18 | 22 |
| Alkaline phosphatase (AP) (U/l) | 1,051 | 814 | 1,113 |
| Bilirubin (µmol/l) | 144 | 107 | 85 |
| Leukocytes (Tsd/µl) | 6.8 | 5.4 | 6.5 |
| Urea (mmol/l) | 11.5 | 22.3 | 9.8 |
| International Normalised Ratio (INR) | 1.2 | 1.2 | 1.1 |
| Cell number of polymorphonuclear cells (PMN/µl) | 2,437 | 481 | 435 |
| Creatinine (µmmol/l) | 235 | 394 | 260 |
| Gamma-glutamyl transpeptidase (U/l) | 341 | 255 | 448 |

**Notes:**

".." indicates missing data.

CRP: C-reactive protein, PCT: Procalcitonin, AST: Aspartate transaminase, ALT: Alanine transaminase, AP: Alkaline phosphatase, INR: International Normalized Ratio, PMN: Polymorphonuclear cells.

### S3 Table. Antibiotic susceptibility profiles of the *E. coli* isolate

| Antibiotic | Isolate | MIC(μg/mL) |
|---|---|---|
| Ampicillin | R | >=32.0 |
| Ampicillin-Sulbactam | R | 16.0 |
| Piperacillin-Tazobactam | S | 8.0 |
| Cefuroxime | R | 32.0 |
| Cefuroxime-Axetil | R | 32.0 |
| Cefpodoxime | R | 2.0 |
| Ceftriaxone | S | <=1.0 |
| Cefotaxime | S | <=1.0 |
| Ceftazidime | S | <=1.0 |
| Gentamicin | S | <=1.0 |
| Tigecycline | S | <=1.0 |
| Levofloxacin | R | >=8.0 |
| Ciprofloxacin | R | >=4.0 |
| Moxifloxacin | R | >=8.0 |
| Meropenem | S | <=0.25 |
| Ertapenem | S | <=0.5 |
| Cotrimoxazole | R | >=320.0 |

**Notes:**

R: Resistant, S: Susceptible.

MIC: Minimum Inhibitory Concentration.

Antibiotics were tested using standard methods, and susceptibility was determined based on CLSI guidelines.

### S4 Table. *P. yeei* sequences containing pathogenicity associated genes

| Gene | Accession | Category | Count |
|---|---|---|---|
| Beta-lactamase | IPR040239 | Antibiotic resistance | 1 |
| Multidrug resistance protein EmrB | IPR004638 | Antibiotic resistance | 2 |
| Fosfomycin resistance protein AbaF | IPR036259 | Antibiotic resistance | 2 |
| Ribose import ATP-binding protein RbsA | IPR003439 | Secretion | 4 |
| Undecaprenyl-diphosphatase | IPR003824 | Antibiotic resistance | 10 |
| ZnMc domain-containing protein | IPR034033 | Induce inflammation | 1 |
| ZnMc domain-containing protein (Fragment) | IPR034033 | Induce inflammation | 1 |

**Notes:**

Counts represent the number of occurrences identified for each gene in the sequencing data.

**S5 Table. Summary of published case reports for peritonitis patients with demonstrated *Paracoccus yeei* infections.** The first row includes the susceptibility test results for the *E. coli* isolated from the patient in this study.

| Case studies | Age/ Sex | Admition department | Medical diagnosis | Clinical manifestation | Blood analysis | | | | | Peritoneal fluid analysis | | Diagnostic method | | | Treatment | Outcome |
|---|---|---|---|---|---|---|---|---|---|---|---|---|---|---|---|---|
| | | | | | **WBC** | **Alkaline Phosphatase (AP), Alanine Aminotransferase (ALT), Aspartate Aminotransferase (ASP)** | **Bilirubin** | **creatinine** | **CRP** | **WBC** | **PMN** | **Culture+time** | **Prove** | **Susceptibility test** | | |
| 1 (Our case study) | 38 year old Male | Hepatology department | Decompensated Liver Cirrhosis CHILD C (11 P.), Type 3 Diabetes in Cystic Fibrosis | Deterioration in general condition and increasing ascites | 6,800/ µl | 1,051 IU/L, 24 IU/L, 38 IU/L | 144 µmol/l | 235 µmmol/l | 229.6 mg/l | | 2,437 | Blood culture: No, Peritoneal fluid: E. coli | Shotgun metagenomics Nanopore sequencing | Susceptibility to Piperacillin/Tazobactam but resistant to Ampicillin/ Sulbactam, Cefuroxim, Cefpodoxim, Tigecyclin, Levofloxacin, Ciprofloxacin, Moxifloxacin, Cotrimoxazol | Piperacillin/Tazobactam, meropenem, Fluconazol | Cured, Liver transplantation |
| 2 | 50 year old Female | Emergency department | End stage renal disease, APD | Pain with anorexia, diarrhea, and shivering | 9,190/ µl | | | | 47.7 mg/l | 2,111/µl | 1,598 49% | Blood culture: No, Peritoneal fluid: Bactec Plus Aerobic | VITEK™ 2 GN (BioMérieux), MALDI-TOF, 16S rRNA | Susceptibility to ampicillin and amoxicillin-clavulanate | intraperitoneal (IP) vancomycin (75 mg/L) and amikacin (12 mg/L), amoxicillin 150 mg IP per liter of dialysate. Antibiotic was administrated on CAPD, amoxicillin 250 mg per bags 3 times on a day at the beginning. | Cured |
| 3 | 51 year old Male | | diabetic kidney disease, APD, ESDR | no pain, cloudy effluent | | | | | | 4,390/µl | 84% | On day 3 of treatment, Peritoneal fluid culture in conventional culture medium become positive | | MIC: piperacillin, ≤2 mg/L; sulbactam/ampicillin, ≤2 mg/L; ceftazidime, 2 mg/L; cefepime, 1 mg/L; ceftriaxone, ≤1 mg/L; cefozopran, 2 mg/; meropenem, ≤0.5 mg/L; aztreonam, 16 mg/L; minocycline, ≤1 mg/L; gentamicin, ≤1 mg/L; amikacin, ≤4 mg/L; ciprofloxacin, ≤0.25 mg/L; levofloxacin, ≤0.5 mg/L; and trimethoprim/sulfamethoxazole, 1 mg/L | vancomycin and ceftriaxone | Cured |

| | | | | | | | | | | | | | | | | |
|---|---|---|---|---|---|---|---|---|---|---|---|---|---|---|---|---|
| 4 | 56 year old Male | Emergency department | Hepatocellular carcinoma, type 2 diabetes mellitus | Fever, Joint pain, dyspnea | | | | | | | | Blood culture: aerobic bottle after 41h, subculture in blood agar and chocolate agar 48h | MALDI-TOF, 16SrRNA | MIC: sensitivity for piperacillin-tazobactam: ≤8 g/mL, ceftazidime: ≤1 g/mL, cefepime: ≤1 g/mL, aztreonam: 4 g/mL, meropenem: ≤1 g/mL, amikacin ≤8 g/mL and ciprofloxacin ≤0.5 g/mL. Although others have reported strains resistant to ciprofloxacin or to third-generation cephalosporins, this strain was sensitive to these antibiotics. | meropenem and ciprofloxacin | Cured |
| 5 | 81 year old Female | Peritoneal dialysis (PD unit) | Renal failure secondary to type-2 diabetes mellitus, continuous ambulatory peritoneal dialysis (CAPD) | Cloudy effluent<br>No fever<br>No abdominal pain | 8,900/µl | | | | 10.9mg/l | 105/µl | | Peritoneal fluid: Columbia blood agar (CBA) and chocolate agar 48 h | MALDI-TOF, 16SrRNA | MIC: sensitivity for amikacin, gentamicin, tobramycin, piperacillin/ tazobactam, imipenem and meropenem. Intermediate for Aztreonam, ceftazidime and ciprofloxacin | gentamicin 80 mg and vancomycin 1 g | Cured |
| 6 | 42 year old Male | | Decompensated cirrhosis from genotype 1a (G1a) hepatitis C (HCV) | Sepsis, Lymphedema | | 99 IU/L, 31 IU/L, 86 IU/L | 23 mg/dL | | | | | Blood: aerobic bottle+chocolate agar 2days+2days | MALDI-TOF, 16SrRNA | | piperacillin-tazobactam 3.375 mg every 6 hours, metronidazole 500 mg IV every 8 hours, and vancomycin 125 mg, PO every 6 hours before identification and ceftazidime 1 g every 12 hours after diagnosis | Death |
| 7 | 46 year old Female | Dialysis unit | polycystic kidney disease | No fever, slight abdominal discomfort | | | | | | 790/µl | 75% | Peritoneal fluid: in blood culture positive after 48h | 16S rRNA | | intraperitoneal vancomycin and ceftazidime | Cured |
| 8 | 72 year old Male | | Diabetes, APD | sharp abdominal pain | 6,700/µl | | | | | 350/µl | 32% | Peritoneal fluid culture after 14days | | | IP vancomycin 2 g (as he was previously colonized with MRSA) and gentamicin 50 mg | Cured |

| | | | | | | | | | | | | | | | | |
|---|---|---|---|---|---|---|---|---|---|---|---|---|---|---|---|---|
| 9 | 25 year old Male | Nephrology Unit | APD peritonitis | Apyretic, Abdominal pain | 5,000/ µl | | | | 63.0m g/l | 315/µl | 75% | Peritoneal fluid: 48h brain-heart medium+18h sub culturing in blood agar and chocolate agar | 16S rRNA sequencing(1323bp), VITEK 2 GN | MIC: ampicillin 0.064 mg/l, amoxicillin—clavulanate 0.064 mg/l, ticarcillin 0.25 mg/l, ticarcillin—clavulanate 0.25 mg/l, piperacillin 8 mg/l, piperacillin—tazobactam 8 mg/l, cephalothin 4 mg/ l, cefotaxime 1 mg/l, ceftazidime 4 mg/l, cefepime 2 mg/l, aztreonam 32 mg/l, imipenem 0.125 mg/l, tobramycin 0.5 mg/l, amikacin 0.5 mg/l, gentamicin 0.5 mg/l, and ciprofloxacin 1 mg/l. | IP piperacillin +cephalothin | Cured |
